# Plant cold acclimation and its impact on sensitivity of carbohydrate metabolism

**DOI:** 10.1101/2024.06.04.597423

**Authors:** Stephan O. Adler, Anastasia Kitashova, Ana Bulović, Thomas Nägele, Edda Klipp

## Abstract

The ability to acclimate to changing environmental conditions is essential for the fitness and survival of plants. Not only are seasonal differences challenging for plants growing in different habitats but, facing climate change, the likelihood of encountering extreme weather events increases. In order to better assess and respond to associated future challenges and risks it is important to understand the processes happening during acclimation. Previous studies of acclimation processes of *Arabidopsis thaliana* to changes in temperature and light conditions have revealed a multigenic trait comprising and affecting multiple layers of molecular organization. Here, a combination of experimental and computational methods was applied to study the effects of changing light intensities during cold acclimation on the central carbon metabolism of *Arabidopsis thaliana* leaves. Mathematical modeling, simulation and sensitivity analysis predicted an important role of hexose phosphate balance for stabilization of photosynthetic CO_2_ fixation. Experimental validation revealed a profound effect of temperature on the sensitivity of carbohydrate metabolism.

## 1. Introduction

Dynamic environmental factors affect plant metabolism, growth, and development. Due to their sessile lifestyle, all plants have developed mechanisms to respond to varying magnitudes and time scales of changes in light intensity, temperature, or humidity. The reversible adjustment of metabolism to such factors is termed acclimation and comprises various responses on a molecular and physiological level (Kleine et al. 2021).

Many plant species, e.g., *Arabidopsis thaliana* (Seydel et al. 2022), *Zea mays* (Kingston-Smith et al. 1999; Pandolfi et al. 2016) and *Nicotiana tabacum* (Vranová et al. 2002), have been studied to reveal insights into the processes happening during acclimation to e.g. cold, heat, fluctuating or high light. When exposed to temperature changes, besides the need for protection from freezing or adverse heat, all vital chemical processes need to be maintained.

Because the velocity and balance of involved chemical reactions is differently affected by temperature, these processes are likely to become unbalanced. This effect needs to be compensated. The compensation can happen actively, e.g. through changes in gene expression or molecular modifications, or passively, e.g. through the architecture of the reaction network and the physical properties of its components (Rouff et al. 2007). Besides having a compensatory nature, the reaction network can offer potential for regulation (Adler et al. 2020). Taken together, these processes form a complex system of intertwined mechanisms that is difficult to entangle. Following approaches that combine different experimental procedures with mathematical modeling has shown to be a fruitful strategy to address complex problems like this.

During cold acclimation of *Arabidopsis thaliana*, a reprogramming of several metabolic processes has been described. This includes the accumulation of soluble sugars, such as sucrose and raffinose, which potentially act as cryoprotectants and stabilize metabolism (Klotke, 2004; Ristic & Ashworth, 1993; Seydel, 2022). This is accompanied by the induction of *BAM3* expression (Sicher 2011; Kaplan 2004; Monroe 2014), an amylase that is involved in starch breakdown, leading to a decrease of starch amounts during the first hours of acclimation and supporting the accumulation of soluble sugars (Yano et al. 2005). Later during acclimation, starch accumulates (Klotke et al. 2004). Additionally, the activity of TCA cycle enzymes and oxidative respiration is reduced, resulting in a decrease of respiration and energy production (Fürtauer 2019). The activity of enzymes involved in photosynthesis is inhibited, causing a decrease in carbon fixation and the accumulation of reducing equivalents, such as NADPH, which are used for biosynthetic processes (Hannah 2005; Savitch 2001; Fürtauer 2019). Alongside these processes within the primary metabolism, also secondary metabolism undergoes several rearrangements. Here, the accumulation of anthocyanins has been observed (Korn et al 2008). Anthocyanin synthesis is a downstream branch of flavonoid synthesis which branches from the shikimate pathway (Austin & Noel, 2003). It has been demonstrated that this accumulation process is vital for cold protection (Chalker-Scott 1999; Schulz 2015), but while the full scope of anthocyanin function is not completely clear, there is strong evidence for photoprotection, protection from stress-induced ROS and osmoregulation (Agati 2021; Chalker-Scott 2002; Kaur 2023). Since both, primary and secondary metabolism are differentially regulated during cold acclimation and play vital roles for the plants’ survival, there is need for precise regulation of carbon distribution within the different domains of metabolism, that might include regulatory interactions.

Different genotypes of *Arabidopsis thaliana* exhibit different characteristics in their reprogramming of metabolism that are related to their sensitivity to cold. There are, for example, indications for differences of interconversion rates between hexoses and sucrose or of export rates to sink organs (Nägele 2010). Mutants like *bam3, pgm1, chs* and *f3h* are deficient in processes directly involved in acclimation. The *bam3* mutant is deficient in amylase activity that is upregulated in cold conditions. This leads to insufficient retrieval of carbon from starch. The *pgm1* mutant, in contrast, is starch deficient because it lacks phosphoglucomutase that catalyzes an initial step of starch synthesis (Stitt & Zeemann 2012). While these mutations are starch related and, therefore, correspond to primary metabolism the *chs* and *f3h* mutations are related to secondary metabolism and are defective in flavonoid biosynthesis. Chalcone synthase (CHS) catalyzes the diversion of phenylpropanoid tetraketide from the shikimate pathway, the initial step of flavonoid biosynthesis. In a later step, F3H and F3’H (flavanone 3-hydroxylase and flavanone 3’-hydroxylase) catalyze the conversion of naringenin, a precursor of a subgroup of flavonols including anthocyanins, into dihydroflavonol (Owens 2008; Shi & Xie 2014).

Despite the numerous successful studies using different experimental and computational approaches, many aspects of the complex and dynamic mechanisms and processes during cold acclimation are still elusive. In an earlier study (Kitashova et al. 2022), the natural accession Columbia-0, together with the mentioned four mutant genotypes *bam3, pgm1, chs* and *f3h* were used to investigate regulatory interactions between primary and secondary metabolism during acclimation. Data on metabolite levels, enzyme activities and photosynthetic carbon uptake was obtained and used to calibrate a condensed mathematical model, comprising core elements of the central carbon metabolism of plant leaves. In this way, evidence for dynamic carbon partitioning between starch, sucrose and anthocyanin metabolism, including a unidirectional signaling link between starch and flavonoid metabolism and a central role of sucrose could be identified.

In the present study, we analyzed this model to investigate the effects of variations in light intensity on the five genotypes of *Arabidopsis thaliana* during and after two weeks of cold acclimation. We re-estimated the model parameters with literature-based constraints for coherent enzyme specific values across the different genotypes and investigated the sensitivity towards dynamics of net photosynthetic CO_2_ assimilation rates after different periods of high light exposure at 4°C *in vivo* and *in silico*. A comparison of the dynamics showed a good agreement between experiments and simulations. This enabled for broader analysis of these dynamics through further simulations and the introduction of a sensitivity measure (sensitivity score). In combination with the analysis of control coefficients within the system, this approach indicates that temperature has a stronger impact on the overall sensitivity of central carbon metabolism than the different mutations and that there is likely one or more compensation or rescue mechanisms shared by primary and secondary metabolism.

## 2. Methods

### 2.1. The model

#### 2.1.1. Model overview

This work is based on the model presented before (Kitashova et al. 2022). In this study, different genotypes (wild type: Col-0; mutants: *chs, f3h, bam3, pgm1*) of *Arabidopsis thaliana* were exposed to 4°C over the course of 14 days. On days 0, 1, 3, 7 and 14, samples were taken and net photosynthesis (nps), levels of starch, soluble carbohydrates, hexose phosphates, anthocyanins, and organic acids, as well as enzyme activities were quantified. The experimental data was then used to estimate parameters for simulation of the central carbon metabolism and its interface with flavonoid biosynthesis.

The basic model comprises 5 metabolites and 12 reactions and is described by a set of ordinary differential equations (ODEs) given in Tab. 1. A scheme of the model is shown in Figure 1.

**Figure 1:**
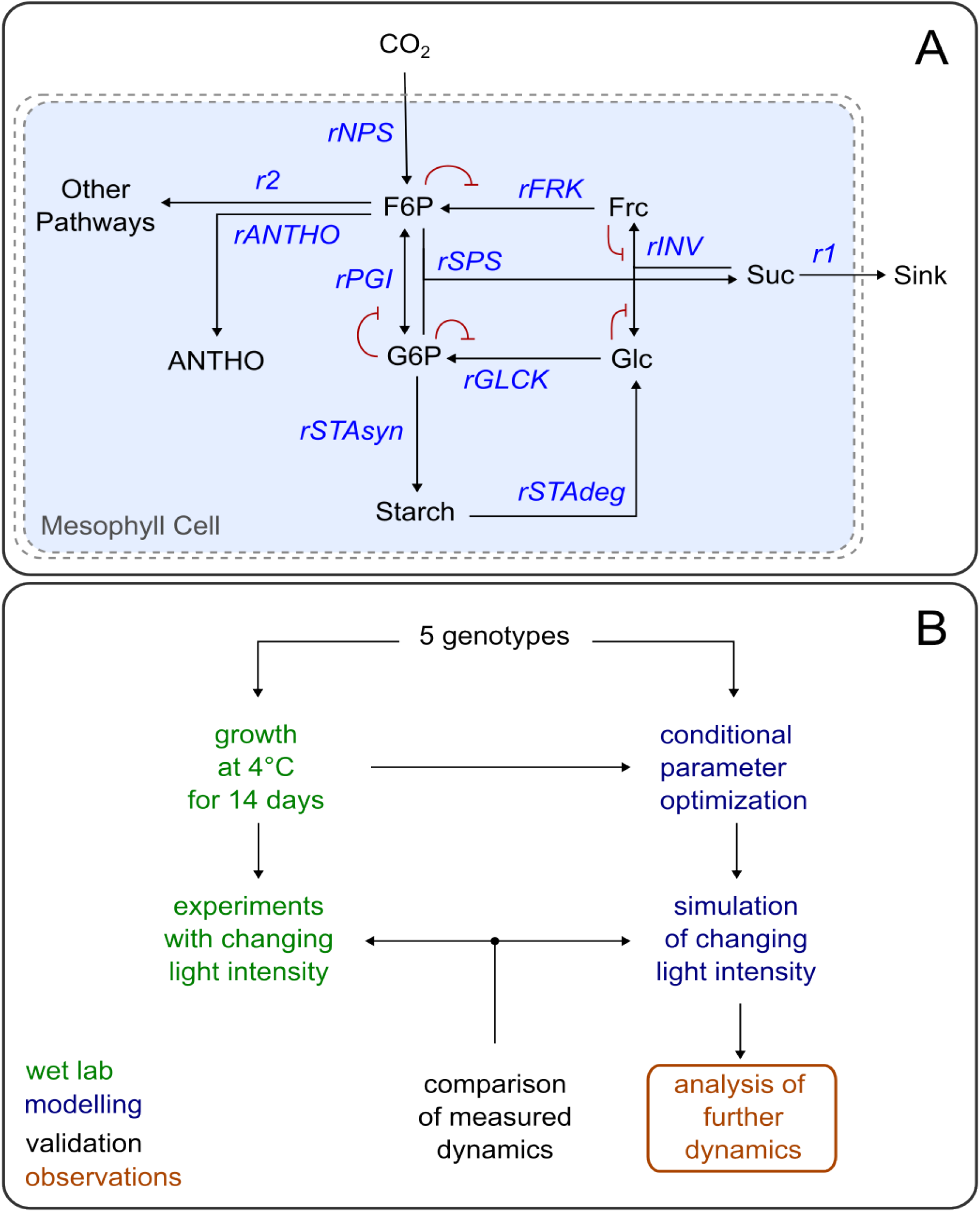
Model scheme and workflow. **A**: The model consists of 5 main metabolites: F6P, G6P, Frc, Glc and Suc. These metabolites take part in various enzymatic reactions (*rSPS, rPGI, rFRCK, rGLCK, rINV*) or biological processes (*rNPS, rSTAsyn, rSTAdeg, rANTHO*). These are depicted as arrows. There are two export fluxes, r1 to sink organs and r2 to other pathways of the central carbon metabolism in the mesophyll cell. The red lines indicate negative feedback regulation by metabolites on reaction fluxes. **B**: This flowchart provides an overview on the workflow and steps in this study. It highlights the experimental (green) and modeling (blue) part and how these are connected. Observations and validation are represented in orange and black, respectively.

In the model, fructose 6-phosphate (F6P) is assumed to be the main product of photosynthesis available for further processes. It is in an equilibrium with glucose 6-phosphate (G6P), that is defined by phosphoglucoisomerase (PGI) activity. F6P and G6P together can be converted into sucrose (Suc) by sucrose phosphate synthase (SPS), while only a fraction of G6P is available for this reaction. Here, UDP-glucose as an intermediate metabolite was omitted for simplification, with the assumption that about 20-50% of G6P are converted before entering the SPS reaction. Suc is then directly exported to sink organs. Additionally, F6P is converted into anthocyanins (ANTHO) and diverted to the second export flux (r2) while G6P is converted into starch in a single process (rSTAsyn) lumping the stepwise synthesis. The invertase reaction (rINV), breaking down Suc into fructose (Frc) and glucose (Glc), together with the fructokinase and glucokinase reactions (rFRCK, rGLCK), phosphorylating Frc and Glc and resulting in the recovery of F6P and G6P, form a central loop that enables responsive balancing of metabolites.

#### 2.1.2. Model equations

##### Differential equation system

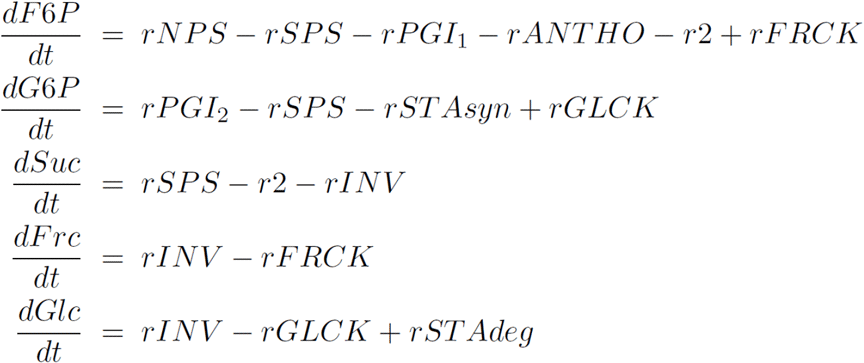

##### Rate equations

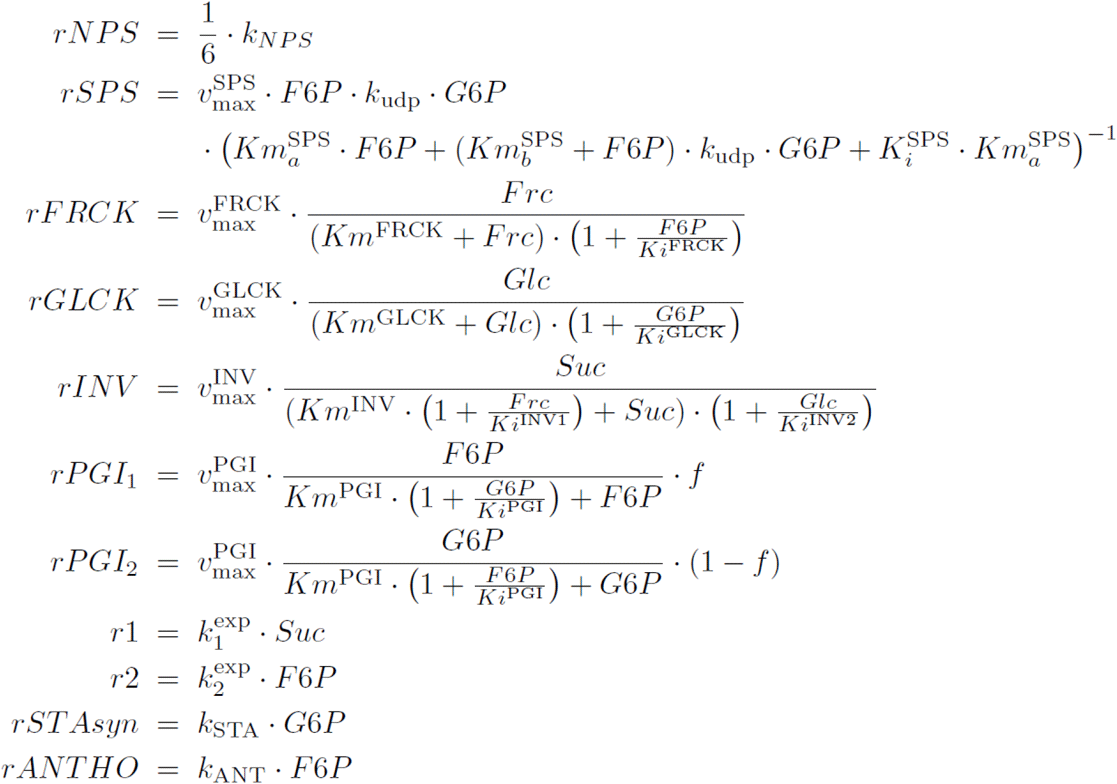

#### 2.1.3. Model refinement and parameter re-estimation

Matching the composition of data presented in Kitashova et al. 2022, the model parameters were optimized to fulfill a steady state assumption for each genotype and time point pair, resulting in 25 conditional model realizations (parameterizations), representing different stages of acclimation for the different genotypes. While keeping the original structure

All corresponding estimated K_m_ and K_i_ values are within a range of one order of magnitude between each set. It is reasonable to assume the possibility of changes in K_m_ values over time, especially during a process of metabolic rearrangement like acclimation due to, for example, post translational modifications such as phosphorylation (Huber and Huber 1996; Ashihara, Crozier, and Komamine 2010; Boex-Fontvieille et al. 2014).

This way, we were able to identify a new refined set of parameters for each combination of genotype and time point, sharing a common, close range of K_m_ and K_i_ values. In the context of parameterization, it is also important to mention of the model, here, we applied a different parametrization strategy by restricting all K_m_ and K_i_ values to common bounds for all genotypes and time points. Additionally, these bounds had to meet conditions derived from literature (Preiser et al. 2020; Dai et al. 1995; Lou et al. 2007; Qi et al. 2007; Xiang et al. 2011):

- K_mFRCK_ > K_mGLCK_
- K_mINV_ similar or close to K_mFRCK_
- K_mFRCK_ > K_mPGI_ > K_mFRCK_

that, while the model considers starch degradation, the degradation rate was set to 0 for the 25 conditional parameterizations, since they represent steady state conditions during midday and most starch degradation usually happens during the night under ambient temperature / control condition. But, as the starch production rates are estimated in a net balance over several days as described in Kitashova et al. 2022, relative changes of mean starch production and degradation are summarized in the production rate parameter *k*_*STA*_. The model with the systematically revised parameters was then subjected to a sensitivity analysis.

### 2.2. Simulations and analyses

For all simulations the free software tool Tellurium (version 2.2.7, Choi 2018) was used, utilizing the CVODE integrator to solve the set of ODEs. To perform metabolic control analysis, methods of libroadrunner (Somogyi et al., 2015; Welsh et al., 2023) within Tellurium were used. The free Matlab software package Data2Dynamics (Raue 2015) was used for parameter estimation.

### 2.3. Experimental procedures

#### 2.3.1. Plant material and growth conditions

*Arabidopsis thaliana* accession Col-0 and homozygous T-DNA insertion lines *bam3* (beta amylase 3, line SALK_041214, locus AT4G17090), *chs* (chalcone synthase, line SALK_020583C, locus AT5G13930), *f3h* (flavanone 3-hydroxylase, line SALK_113904C, locus AT3G51240), as well as the SNP mutant *pgm1* (plastidial phosphoglucomutase, TAIR stock CS3092, locus AT5G51820) were grown on a 1:1 mixture of GS90 soil and vermiculite in a climate chamber under controlled conditions (8h/16h day/night; 100 μmol m^−2^ s^−1^ ; 22°C/16°C; 60% relative humidity). Following two weeks of further growth under long day conditions (16h/8h day/night; 100 μmol m^−2^ s^−1^ ; 22°C/16°C), plants were shifted to 4°C (16h/8h day/night; 90-100 μmol m^−2^ s^−1^ ; 4°C/4°C). On the 14th day of cold, plants were either (i) sampled after 3h of ambient light; or (ii) exposed to elevated light (240-270 μmol m^−2^ s^−1^ ; 4°C/4°C) and sampled after 3h and 6h of elevated light. Each sample consisted of 1 leaf rosette which was snap-frozen in liquid nitrogen, ground to a fine powder and freeze-dried.

#### 2.3.2. Hexose phosphate quantification

Fructose 6-phosphate (F6P) and glucose 6-phosphate (G6P) amounts were quantified as described before (Gibon et al., 2002) . In brief, F6P and G6P were extracted with 16% (w/v) trichloroacetic acid in diethyl ether and washed with 16% (w/v) trichloroacetic acid in 5 mM EGTA. F6P and G6P amounts were determined in a 2-step cycling assay, catalyzing equimolar interconversion of hexose-phosphates into NADPH + H^+^ , which was photometrically determined in a reaction with thiazolyl blue and phenazine methosulfate at 570 nm.

## 3. Results

In order to investigate the influence of dynamic CO_2_ assimilation rates on the acclimation process of the central carbon metabolism in *Arabidopsis thaliana* leaves, we investigated the presented model in a restricted parameter space. Therefore, we defined a procedure for assessing the dynamic sensitivity to these perturbations and compared its outcome when applied to model simulations or new experimental data. Additionally, we embedded the condensed model into a large-scale metabolic model and compared the resulting fluxes when constrained accordingly for further validation.

Our presented analysis, together with the evaluation of parameter control coefficients, could reveal an important role of hexose phosphate balance, a rearrangement of sensitivities in response to temperature change and potential common metabolic compensation mechanisms.

### 3.1. Amounts of F6P and G6P are differentially affected by mutations in flavonoid and starch metabolism

Hexose phosphate amounts were quantified in different light conditions after cold acclimation to assess their sensitivity changes in net photosynthesis rates. While the median values of F6P ranged from 2 to 6 μmol gDW^-1^ those of G6P ranged up to 26 μmol gDW^-1^, the value distributions of the single measurements shared roughly the same range between 0 and 45 μmol gDW^-1^ (Fig. 2). In *chs*, the amount of F6P increased after 3h of high light while amounts were stabilized in all other genotypes (Fig. 2). Both *pgm1* and *bam3* mutants were significantly affected in G6P metabolism, and *pgm1* accumulated highest amounts across all genotypes. However, between 3h and 6h of high light exposure, G6P amounts decreased again in plants of both *pgm1* and *bam3* which was not observed in Col-0, *chs* and *f3h*.

**Figure 2:**
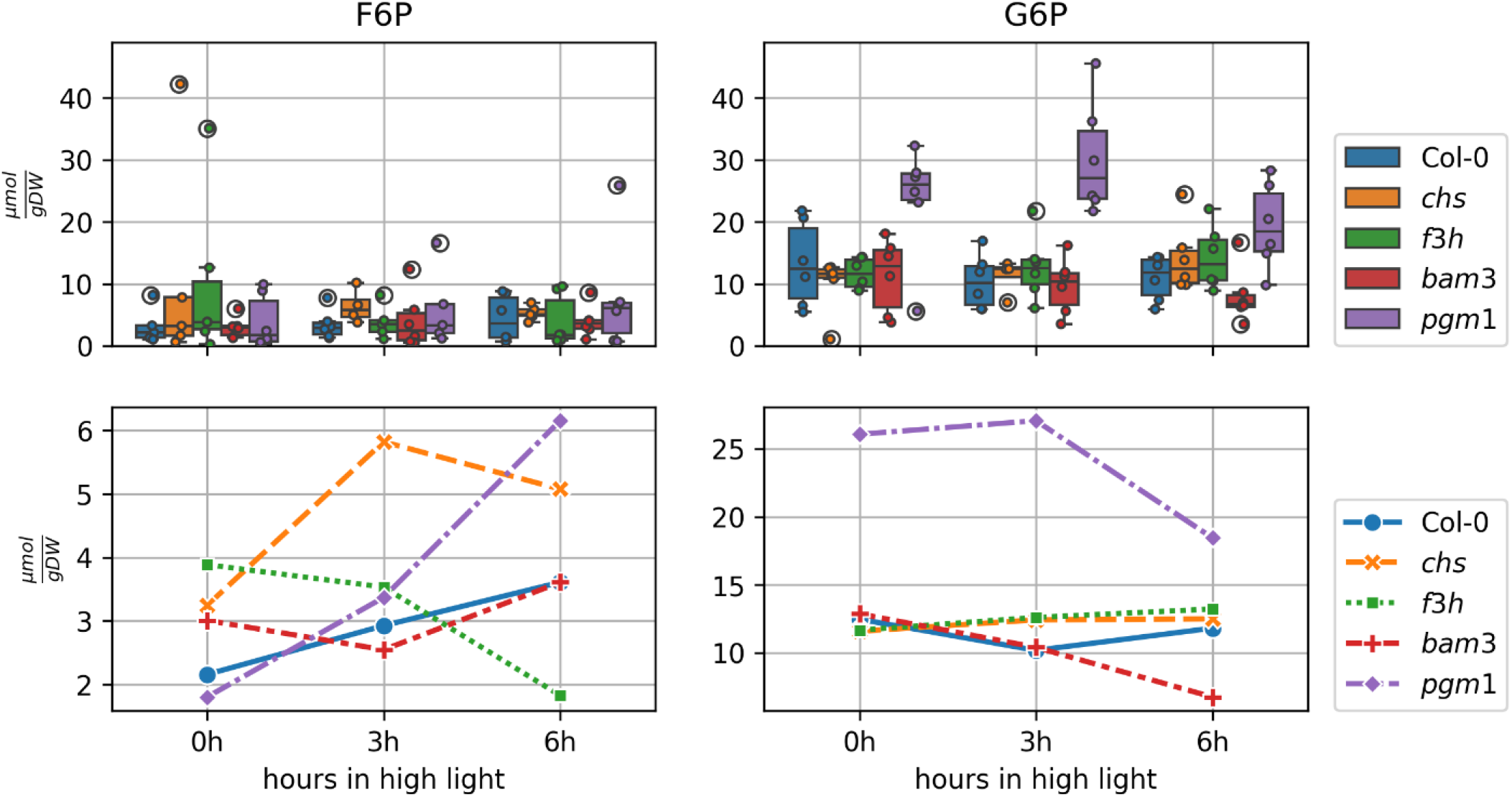
Experimental results. The upper panel illustrates the distributions of hexose concentrations, measured after 0, 3 or 6 hours in high light conditions, as boxplots for each genotype. Here, every single measurement is shown by a color filled dot. The lower panel shows the changes of the measured median over time. The 0h time point is actually represented by samples that were subjected to 3 hours of normal light, as described in the methods section, but is put as 0h time point here for better comparison.

### 3.2. Model validation

#### 3.2.1. Sensitivity analysis

To further assess the impact of acclimation on central carbon metabolism, the different conditional model realizations were subjected to external perturbations, namely changes in net photosynthesis (nps) rates. Under *in vivo* conditions, variations in nps result from naturally

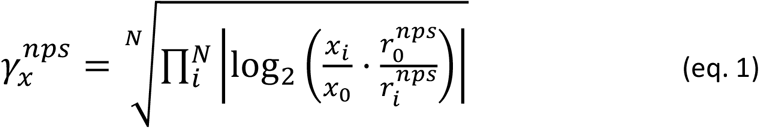

where *γ*_x_^nps^ is the sensitivity score of measure x (metabolite concentration or flux) on perturbations of nps, N is the number of performed perturbations (increase/decrease, 5%, 10%, …), x_0_ is the initial steady state value of measure x and x_i_ the resulting value after changing light. Starting from steady states that were derived from the experimental data, the nps rates in the model were increased or decreased by 5 to 30% in 5% increments and 4h time courses were simulated. The resulting changes in metabolite levels and reaction fluxes were then taken together to compute the sensitivity of the respective condition to these perturbations as perturbation i. The ratio of initial nps rates and rates of perturbation i are represented by r_0_^nps^ and r_i_^nps^, respectively. Figure 3 illustrates this procedure and provides a graphic representation of how well the results match those derived from the experimental data.

**Figure 3:**
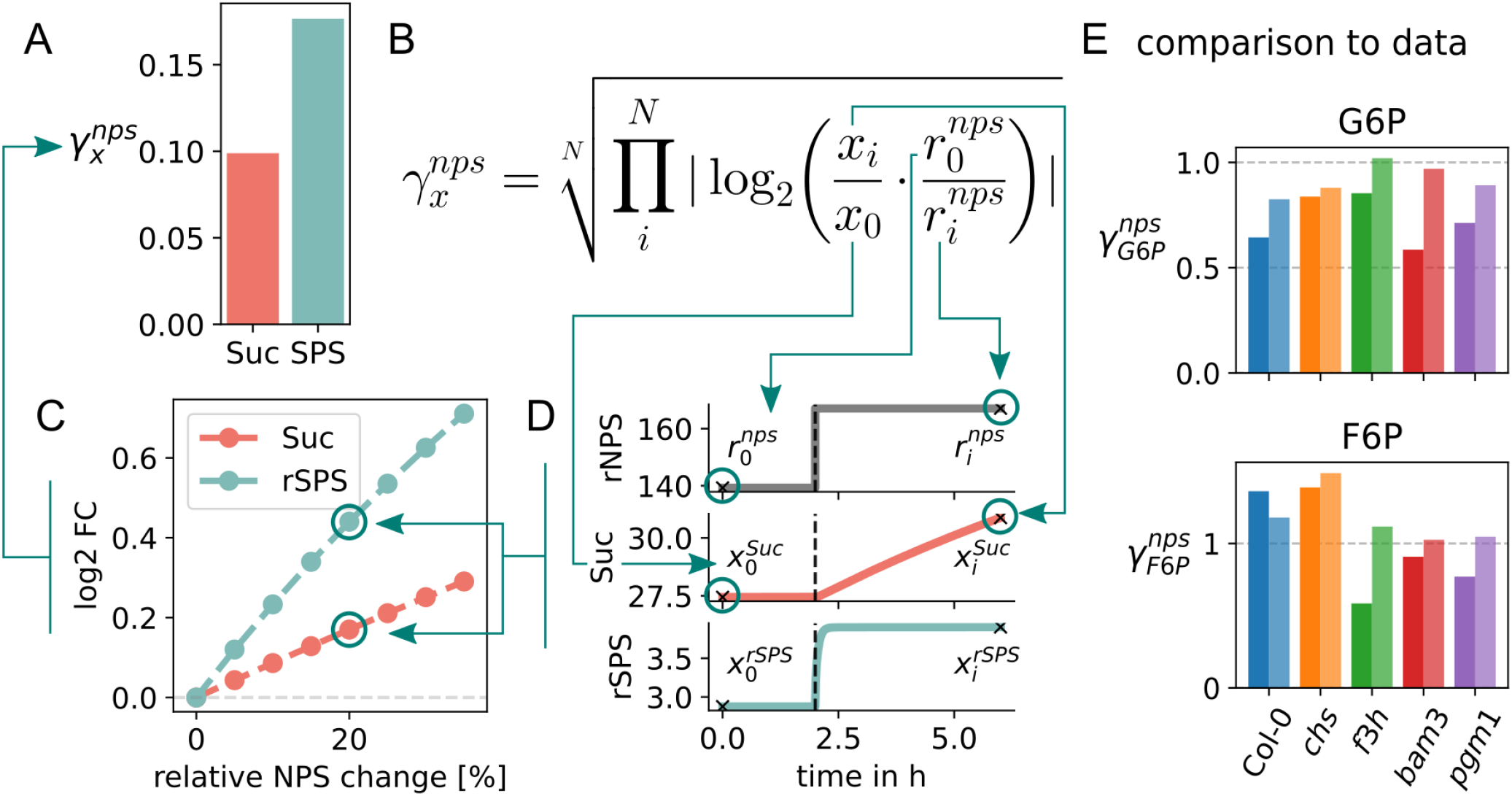
Estimation of sensitivity scores *γ*_x_^nps^ and comparison to experimental data. **A**: Sensitivity scores of sucrose concentration (Suc) and the SPS reaction flux (rSPS) of model Col-0 day 0. **B**: Formula for the sensitivity score *γ*_x_^nps^. **C**: Log_2_ fold-changes of Suc and SPS for computed net photosynthesis (nps) increases. **D**: Exemplary time courses of underlying simulations for a change in nps rate (rNPS) of 20%. Shown are the time courses of rNPS, Suc and rSPS. The vertical black lines indicate the time point of rNPS increase (2h). The “x”s mark the corresponding values for the computation of *γ*_x_^nps^. **E**: Sensitivity scores computed from control experiments (darker bars) are compared to sensitivity scores of simulations (lighter bars).

#### 3.2.2. Comparison of model predictions and experimental data

To evaluate the suitability of the described analysis of sensitivity to variations in nps rates, we compared *γ*_x_^nps^ derived from control experiments to values estimated from results of model simulations mimicking the experimental procedure of a newly performed experiment to specifically test the sensitivity *in vivo*. To this end, *Arabidopsis thaliana* plants from all 5 genotypes were grown at 22°C and transferred to 4°C for 14 days, where they could acclimate as described in Kitashova et al. 2022. On day 14, the plants were either exposed to normal light (100μE) and samples were taken after 3h. Or the plants were transferred to elevated light (240-270μE), which leads to an increase of nps by 20-30%, and samples were taken after 3 and 6h. Concentrations of F6P and G6P were measured for all samples as described in the methods section 2.3.

As a comparison, the model realizations of day 14 were used for each genotype and simulated in the same fashion as the experimental procedure. Here, the shift to high light was simulated as an increase of rNPS by 25%. From the obtained experimental and simulation data *γ*_x_^nps^ values were computed according to eq. 1 with the simple adjustment that, instead of different perturbation intensities, x_i_ represent different durations of exposure (3h and 6h). The concentrations obtained under normal light are the reference values x_0_ and r_0_^nps^. This comparison is shown in Fig. 3 E. We see a very good agreement between experimentally obtained values and simulation results. For each metabolite there is one genotype (*bam3* for G6P and *f3h* for F6P) where the model predictions deviate more than 50% from the data. In summary, the overall agreement verifies a good suitability of the presented sensitivity assessment.

#### 3.2.3. Model fits well into broader metabolic network context

To further validate the model and test its performance, especially considering the degree of condensation, we compared it to a large scale metabolic model.

Because the tackled phenomenon of carbon partitioning into starch and sucrose involves some highly connected metabolites (used in many reactions), we wanted to make sure that our model simplification still fits reasonably well into the broader context of the metabolic network of *A. thaliana*. For this, we compared the flux distribution from the dynamic model proposed here, and the one obtained through a Flux Balance Analysis (FBA) (Orth, Thiele, and Palsson 2010) simulation of a genome-scale metabolic reconstruction of *A. thaliana* known as AraCore (Arnold and Nikoloski 2014).

All comparisons were done for the Col-0 genotype, as the other strains involve either interruptions in the pathways, used to produce biomass precursors (which make the strains non-viable in FBA context), or reduce the efficiency of a cellular process, a subtlety which cannot be considered in a stoichiometry-based modeling framework such as FBA.

We used a number of fluxes from the dynamic model to constrain the AraCore model. The import reaction to F6P was used to constrain the flux through FBPase (both cytosolic and chloroplastic), the major contributor to synthesis of F6P through the Calvin-Benson-Bassham cycle. Additionally, we used the dynamic model fluxes to constrain the fluxes through S6P synthase (S6PS_c), G6P isomerase (PGI_c), and hexokinase (HEXK_c). The lower and upper bound of those reactions were set to 90 and 110% of the flux values obtained in the dynamic model.

We then compared (1) the flux *r2* (flux going out from F6P, see Figure 1) to the flux through F6P transketolase which is a major pathway through which F6P is used for replenishing of the Calvin-Benson-Bassham cycle intermediates, (2) the total flux going out of sucrose in both our dynamic model and the FBA model and (3) the total rate of starch synthesis. One outgoing flux from F6P, which we did not compare to the AraCore prediction, is the production of anthocyanins. The reason for this is that anthocyanins are not explicitly modeled in the condensed dynamic model, and their modeled precursor, shikimate, is not exclusively used for anthocyanin synthesis. Due to this discrepancy, we decided that this comparison would not be meaningful.

When analyzing the results in detail, we see that the major flux depleting F6P (*r2* in dynamic model) matches well (in order of magnitude and exact quantitative value) to the one predicted by AraCore (see Fig. 4 A). Since these are some of the fluxes with highest absolute values, this indicates that the major stoichiometric constraints on the network are well represented in the dynamic model, and that it can reliably be used as a proxy for the global metabolic state of the modeled metabolites.

**Figure 4:**
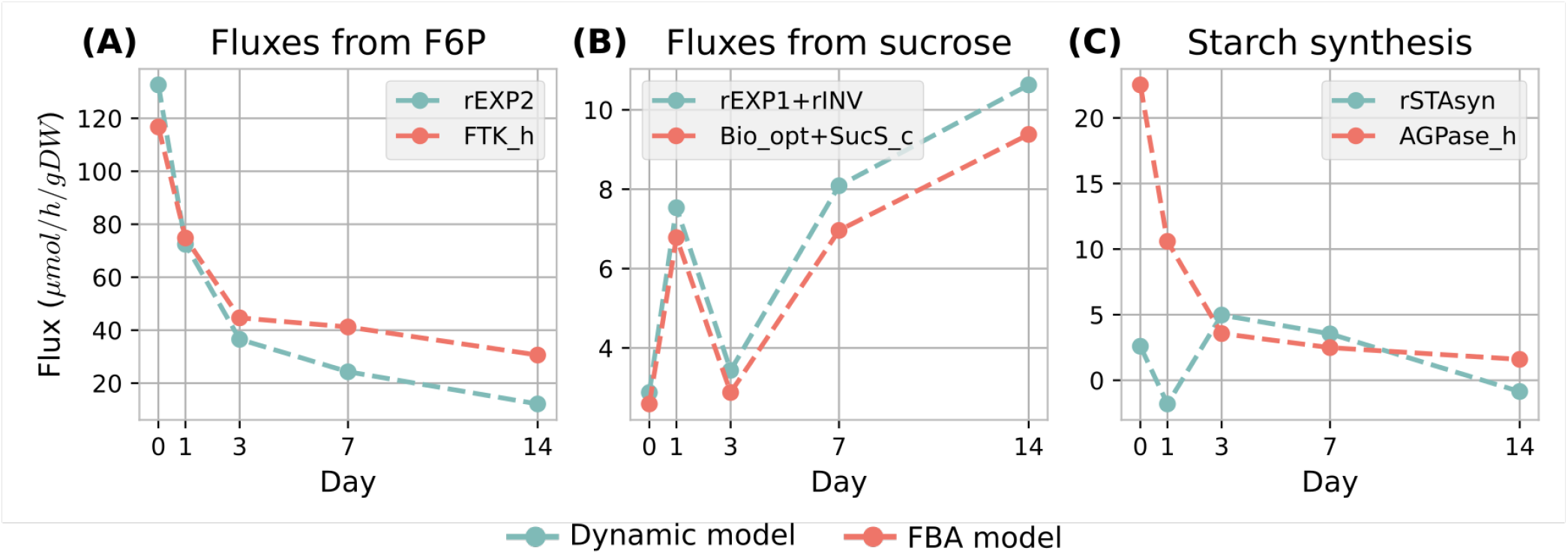
Comparison of flux values between the dynamic model and FBA. **A**: Comparison between *r2* dynamic model flux and the flux through FTK reaction in the chloroplast facilitating replenishment of CBC metabolites. **B**: Comparison of outgoing fluxes of sucrose (invertase and export flux *r1*). **C**: Comparison between starch synthesis fluxes.

Next, the total fluxes going out of sucrose match almost exactly in both models (see Fig. 4 B). However, we have to point out that the values of two individual fluxes that go out of sucrose in the dynamic model, i.e. the export flux *r1* and the flux through the invertase, do not match well to their corresponding fluxes in the FBA model. The predicted export flux of sucrose is much smaller in the FBA model due to the fact that it is represented directly by the biomass reaction. Since the biomass reaction requires many metabolites in a particular stoichiometric relation, the flux through it cannot be easily changed, and is determined not only by sucrose, but by the entire state of the metabolic network. The comparison of starch synthesis does not match as well as the other two comparisons, even if the order of magnitude is not far off, especially after day 3 (see Fig. 4 C). We assume this is because of the way in which the starch synthesis rate was initially calculated from starch measurements, by averaging the difference between starch quantity in the morning between consecutive days over 24 hours (net balance).

Overall, we see a good agreement between the fluxes concerning the modeled metabolites in the dynamic and in the FBA model, giving further confidence in the developed model, the measurements used to calibrate it, and the calibration itself. On this basis, we will subject the model to a thorough analysis of parameter sensitivity in order to understand the role of individual parameters in the acclimation ability of the plants.

### 3.3. Observations and Insights

#### 3.3.1. Hexose phosphate balance has a big impact on the overall carbon distribution

After investigating the plausibility of the model and parameters, first by comparing the sensitivity to variation of nps rates with new *in vivo* data and, second, by comparing the fluxes of our small dynamic model to an established genome-scale FBA model, we also did a classical sensitivity analysis by computing all flux, concentration, and parameter control coefficients to assess the impact of parameter choices on the model behavior. We used the control coefficients in the definition according to

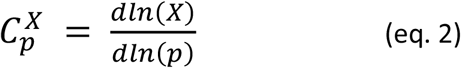

where C^X^_p_ is the control coefficient of flux, concentration or parameter *p* over measure *X* (flux or concentration) (Heinrich & Rapoport 1974; Kacser & Burns 1973). It describes how strong a variation of *p* affects the steady state values of *X*. In this sense, it differs from the approach described before, because it always considers a relaxed system.

The results of this systematic sensitivity analysis are shown in Figure S2. In general, we see a variable influence of most parameters, depending on the given conditions, while no clear trend is apparent. For some parameters, the control coefficients show 2 modes, representing conditions under which the given parameter has an influence or doesn’t have an influence (e.g. due to substrate depletion). The same observations hold for fluxes and concentrations.

Notably, the parameter control coefficient of *f*, the parameter that determines the balance between F6P and G6P at the PGI reaction was significantly (4-5 times) larger than all other control coefficients.

An overview of this analysis is provided in supplementary section 1.

To identify critical parameters and conditions, control coefficients with values larger than 1.5 were investigated more closely. This is the point at which the distribution of control coefficients exhibits a clear drop and a relatively wide tail, meaning that most values are smaller, but a few are considerably larger than 1.5. Interestingly, these values could not clearly be attributed to certain days or genotypes during acclimation, but largely to fructose, glucose and reactions connected to them. This suggests a comparably high sensitivity of this part of central carbon metabolism to perturbations within the system. Since this lack of dependence on genotype or duration of cold exposure was an unexpected result, we decided to investigate whether there might be a sensitivity dependence on temperature instead.

#### 3.3.2. Model sensitivities are more temperature than genotype specific

The investigation of high value control coefficients revealed two additional indications. First, when comparing the frequency of these values between genotypes the values increase from *bam3* along the flavonoid mutants to Col-0 arriving at *pgm1* with the highest frequency. Thus, defects in either starch synthesis or degradation seem to have opposing effects on this centralization of increased sensitivities around Frc and Glc. Second, when comparing the frequency of high value control coefficients between days, day 0 significantly stands out, having a much smaller frequency (10 compared to 35-50). This means that the described sensitivities around Frc and Glc are particular to low temperatures, as they clearly become more pronounced during and after acclimation to 4°C. Both of these observations are illustrated in Figure 5 A, B.

**Figure 5:**
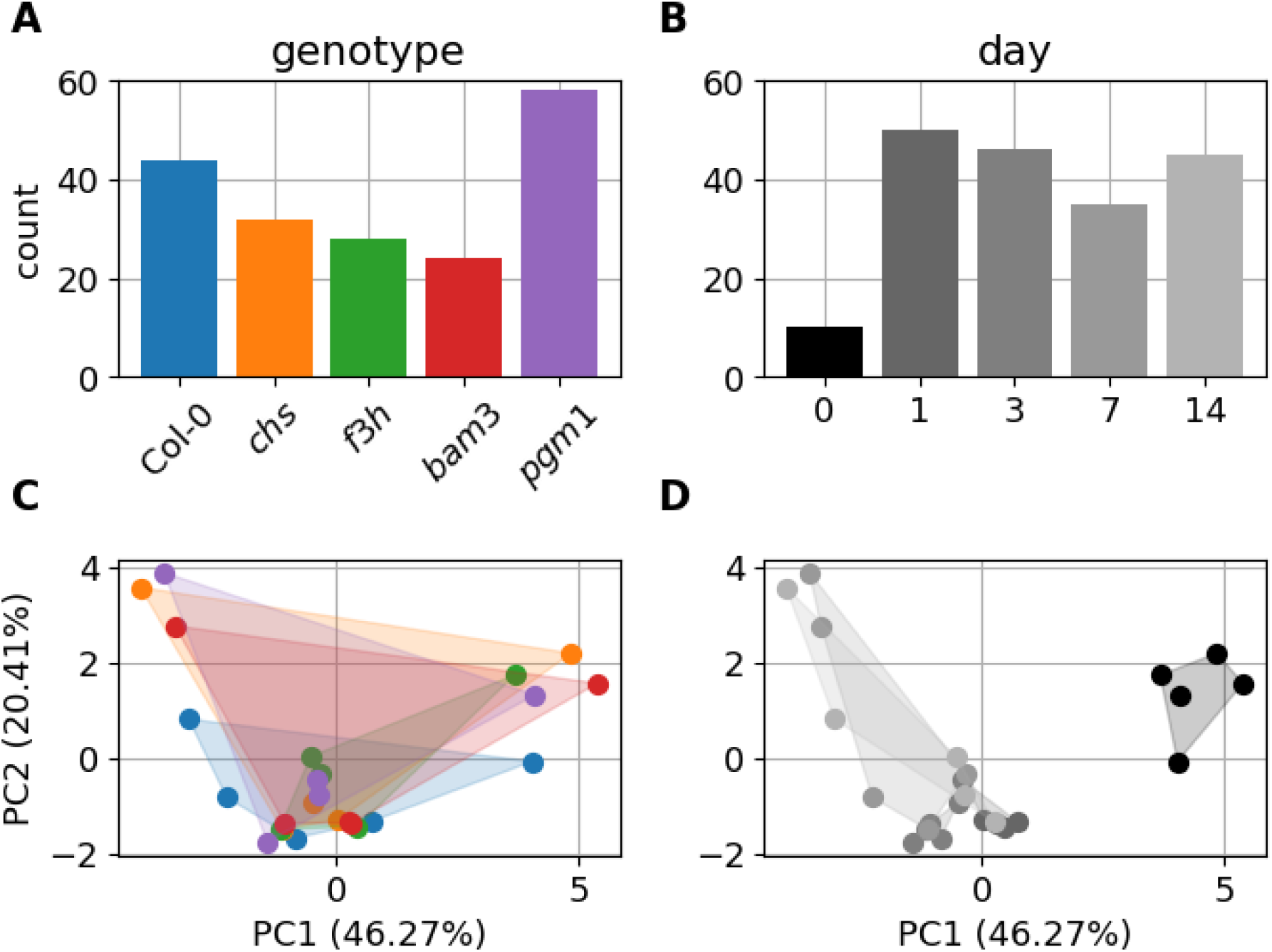
Comparison of model features by genotype or day. The upper panel shows the frequency of parameter control coefficients larger than 1.5 for each genotype (**A**) or day (**B**). The lower panel shows a two-component PCA for the variations in *γ*_x_^nps^. The color attribution of the dots in the lower panels correspond to those in the upper panel (**A** → **C, B** → **D**). The shaded areas in the PCA plots serve a better identifiability of related dots.

To investigate differences in nps variation responses between different genotypes or throughout the course of acclimation, a principal component analysis on *γ*_x_^nps^ was performed. The two components with the highest contribution make up 46.27% (PC1) and 20.41% (PC2) of the differences between the values of all model realizations. The composition of these two components is given in Table 2. The analysis did not show any apparent trend when comparing different genotypes, but interestingly, it exposed an observation that goes along with the described investigation of high value control coefficients. When comparing the days, day 0 is significantly shifted along the PC1 axis, compared to all other days, as illustrated in Figure 5 C, D. This means that the variation of nps rates affects the system in a considerably different manner before and during or after the acclimation process. Since PC1 and PC2, together, account for 66.68% of the variation there are still components PC3 and PC4 adding a considerable amount to the variation with 14.72% and 11.77%, respectively. These components were also investigated, but no clusters or other particularities could be observed. A visualization of these components is provided in the supplementary Fig. S2.

**Table 2.**
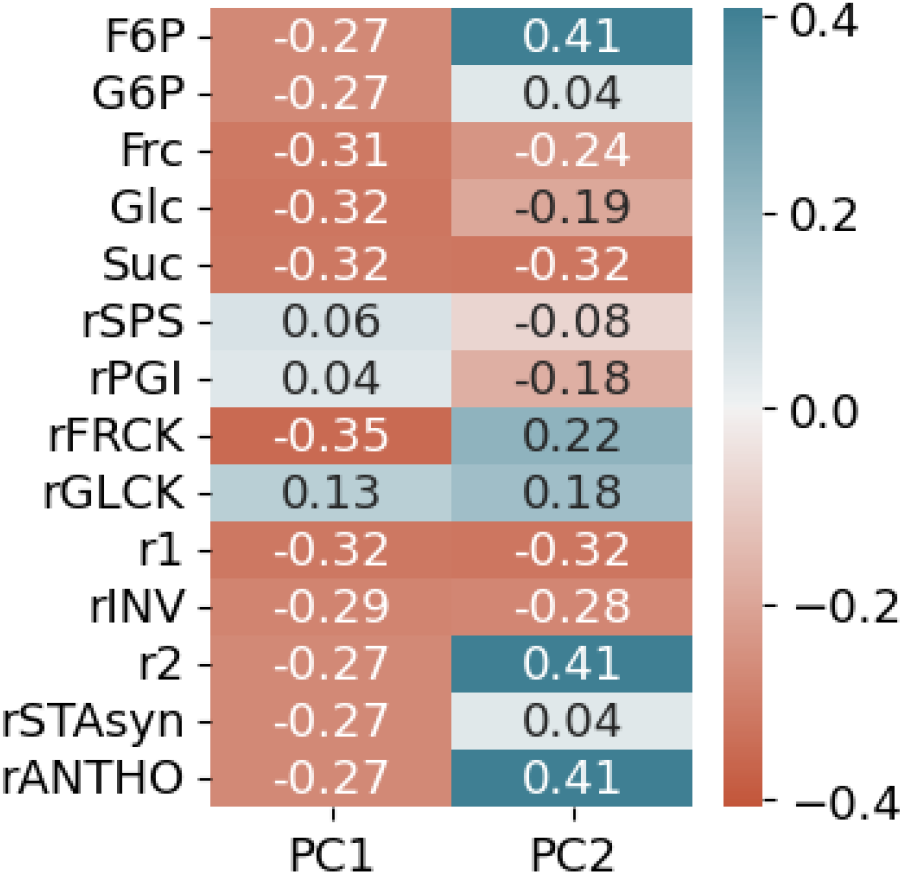
PCA component composition.

#### 3.3.3. Mutants with deficiencies in reactions of the central carbon metabolism exhibit similar deviations from the wild type

An overview of *γ*_x_^nps^ values for all genotypes and days is given in Figure 6. We see that the rFRCK and rGLCK reactions are particularly sensitive to nps rate variations, and also the concentrations of Frc and Glc have relatively high values over all genotypes. Alongside an increase of rFRCK sensitivity over the course of acclimation, also those of F6P, anthocyanin synthesis and r2 increase. While peaking on day 14 for Col-0 and the flavonoid deficient mutants (*chs* and *f3h*), the peak already occurs on day 7 for the mutants with defects in starch metabolism (*bam3* and *pgm1*). Interestingly, the sensitivities of rFRCK, rGLCK, Frc, Glc are prominent compared to values of other measures, but are in a close range over all genotypes. This becomes evident when looking at the heatmaps in the right part of Fig. 6 depicting the log_2_ fold changes of all *γ*_x_^nps^ values compared to the corresponding values of Col-0. The fold changes of rFRCK, rGLCK, Frc, and Glc are always close to zero for all mutants.

**Figure 6:**
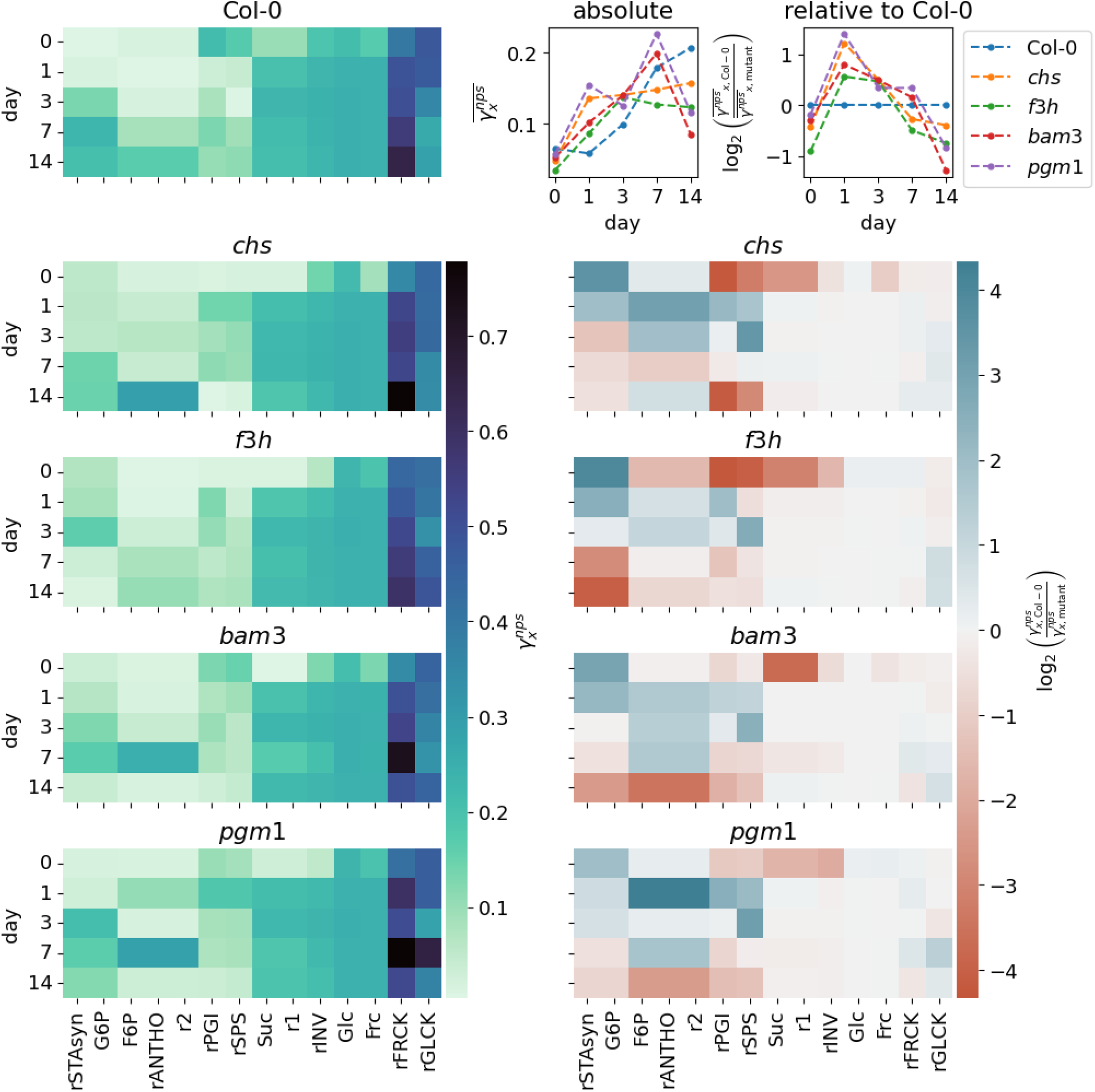
Overview of *γ*_x_^nps^ values. The heatmaps on the left side show the *γ*_x_^nps^ values for every model realization. For each genotype the evolution of values can be followed over days from top to bottom. Each column represents a different measure of the model, that is either a metabolite concentration or a flux. They are not ordered by type, but in the sense of related blocks within the model structure. The heatmaps on the right side show log_2_ fold-changes of these values when compared to the corresponding value of Col-0. The panel in the top right corner illustrates the evolution of the geometric mean of all *γ*_x_^nps^ values per day for each genotype. The left panel shows the absolute values, while the right panel, again, shows the log_2_ fold-changes compared to Col-0.

Examining the other measures, we see a set of divergence patterns that appear to be conserved over all considered mutants. Clearly, there are apparent quantitative differences, but the qualitative behavior is fairly robust. The fold change for G6P and starch synthesis (rSTAsyn) starts at positive values and constantly declines, switching to negative values around day 3 for all mutants. Values of F6P, anthocyanin synthesis (rANTHO) and r2 start from a low or slightly negative fold change, switch to positive values during acclimation and return to a low or negative value after acclimation or, in the case of *chs*, already at day 7. The reaction fluxes rPGI and rSPS follow a similar pattern but return to low or negative values already at day 7 for all mutants. Here, the flavonoid mutants both show more negative values on day 0 than the starch related mutants. Suc, r1 and rINV start with negative fold changes that vanish during and after acclimation.

When following the mean *γ*_x_^nps^ values for each genotype over the course of days, as illustrated in the top right corner of Figure 6, one can notice an overall increase of mean absolute sensitivity from day 0 to day 14. Considering Col-0 as the baseline sensitivity, we again see a similar pattern for all mutants, that is comparable to the behavior of PGI and SPS with a stronger decline at the end, due to the contributions of G6P and starch synthesis.

It is difficult to interpret these observations in terms of defined mechanistic insight. However, the emergence of similar divergence patterns from the wild type indicates that common rescue mechanisms might compensate for the deficiencies of these mutants, even if different parts of the central carbon metabolism are affected.

## 4. Discussion

In this study we present an in-depth analysis of a condensed dynamic model of the core central carbon metabolism of *Arabidopsis thaliana* leaves. It is a model that has already been used to identify a role of limitations in sucrose synthesis in carbon partitioning during acclimation (Kitashova et al. 2022). Since it has been shown that light intensity is of great importance for the cold acclimation process of plants and there are complex interactions between cold and light signaling (Szalai et al. 2018), a dynamic sensitivity analysis was defined, and new experiments were performed to assess its applicability. Also, the model has been refined with additional parameter constraints, according to literature (Preiser et al. 2020; Dai et al. 1995; Lou et al. 2007; Qi et al. 2007; Xiang et al. 2011). Certainly, there are different strategies to calibrate dynamic models. We applied a steady state assumption, because we are considering long term changes (days compared to minutes or hours) and there are measures that were estimated as net balance from the experimental data (starch synthesis) for which this assumption is suitable. In terms of parameter estimation, we tried various procedures. These were: (a) finding the best optimization score for each condition individually without further restrictions, (b) for each condition individually with the restrictions explained in the methods section, (c) for conditions sharing the exact same K_m_/K_i_ values, (d) for each day in a pairwise K_m_/K_i_ fitting procedure. While (c) did not result in any successful optimization, parameter sets were found for (a), (b) and (d). Their performance in comparison to the experimental data was tested and (b) scored the best. Since it also follows reasonable restrictions for the kinetic parameters derived from literature, we are confident that it is best suitable and provides the most reliable predictive power for the presented study. Additionally, it has been embedded into a large-scale metabolic network, resulting in flux values that are in good accordance with those derived from dynamic simulations.

Utilizing the presented sensitivity analysis in combination with a classical approach, using control coefficients, revealed several indications: (i) Classical analysis showed a strong influence of the hexose phosphate balance on overall dynamics. (ii) Both, classical and dynamic sensitivity analysis, demonstrate a more pronounced and discernible impact of temperature on the system’s sensitivity compared to genotype. Classical analysis shows notable differences between different mutants, by an increase of highly sensitive elements due to a lack of starch, but a decrease due to diminished carbon retrieval from starch or defects in flavonoid synthesis. (iii) The dynamic analysis revealed consistent patterns of response deviation from the wild type among mutants with diverse defects in central carbon metabolism, suggesting shared mechanisms to counteract or compensate for these deficiencies.

When looking at the model structure as shown in Figure 1, at first glance, it seems obvious that *f*, the parameter responsible for F6P-G6P balancing in connection to the PGI reaction, can have a strong impact on carbon partitioning, since it defines the fraction of carbon that can flow into the lower right part of the scheme (G6P, Suc, Starch, Glc and Frc) or into the upper left part (F6P, anthocyanins and other pathways), representing primary and secondary metabolism and carbon recycling, respectively. This does not mean it is used as a main regulator of carbon partitioning *in vivo*, but from a modeling point of view, its value has a strong influence on the overall model behavior. Nevertheless, this relationship needs to be balanced because F6P and G6P are both needed for an adequate flux through the rSPS reaction. Interestingly, when looking at the control coefficients of *f* for each system measure, there is a strong positive effect on Glc concentration, but an equally strong negative effect on Frc concentration, which is not immediately obvious by intuition. Here, the negative feedback of G6P plays an important role. Because the F6P/G6P balance emphasizes G6P, but the flux through rSPS is limited, there is an accumulation of G6P, that in turn blocks the activity of glucokinase via a negative feedback, leading to an accumulation of Glc as well. This accumulation of Glc in turn, inhibits invertase activity and consequently leads to a depletion of Frc.

Given the high control coefficients of *f*, the rPGI reaction appears to be the most prominent control point for carbon distribution in the system. However, one needs to pay attention to the fact that the presented model is considerably simplified and condensed and the considered rPGI reaction comprises both, plastid and cytosol reactions. Compartmentalization and subcellular organization has been pointed out to play an important role for the central carbon metabolism and in terms of cold acclimation (Hoermiller et al. 2017; Lundmark 2006). It is an appropriate assumption that not only the ratio of hexose phosphates, but also their subcellular proportions are of great importance for the coordination of carbon partitioning. An extension of this study could focus on exploring this aspect. Nonetheless, these observations underline the critical importance of F6P/G6P balancing and a fine-tuned interplay of related reactions and processes. This notion is in agreement with the indications in Kitashova et al. 2022, as the rSPS reaction directly affects the balance of these hexoses.

Additionally, the new experimental data contains interesting insight connected to F6P/G6P balancing, by revealing that, for starch related mutants, the qualitative changes of F6P concentrations under high light and after acclimation are much closer to those of the wild type, than those of flavonoid related mutants. They increase for longer exposure while those of flavonoid related mutants decrease. With G6P, it is the other way around. For the wild type and flavonoid related mutants G6P concentrations are fairly stable, while those of starch related mutants decrease. Since flavonoid synthesis depends on F6P and starch synthesis depends on G6P, these qualitative deviations from wild type behavior directly indicate the affected parts of central carbon metabolism for each category of mutants.

Although (ii), the more pronounced and discernible impact of temperature, and (iii), the consistent differences to Col-0 in response patterns, are two separate observations, they fit into a coherent picture. Considering a more exposed alteration of the systems sensitivity through a temperature shift compared to the metabolic defects, suggests a successful compensation mechanism for the different mutants and it has been shown that there are mechanisms that can compensate gene knockouts in *Arabidopsis thaliana* (Buzduga 2018). Shared, or partially shared, compensation mechanisms for different deficiencies would be beneficial for the plant, because it would not require a specifically tailored response for all different kinds of limitations, enabling a more economical use of resources. The observed alterations in response patterns hint to a general or shared mechanism like this because the observed patterns are always of the same quality. If different compensation programs were at play, one would expect more apparent differences between the responses, at least between starch related and flavonoid related mutations, like more pronounced or opposing sensitivity changes for some reactions or metabolites. It has been observed that most metabolic heat shock responses are also responses to cold shock (Kaplan et al. 2004), demonstrating the existence of common stress response mechanisms in the context of central carbon metabolism.

Even though these similarities are remarkable there are noticeable differences. When looking at the temporal evolution of the aggregated sensitivities in Fig. 6, we see that the starch related mutants show the highest deflection and overall deviation from Col-0. Also, the aggregated sensitivity of Col-0 increases up until day 14. While the timecourses of the flavonoid related mutants seem to saturate, the values of the starch related mutants drastically decrease again between day 7 and 14, indicating that this enhanced sensitivity cannot be maintained throughout the acclimation process by these mutants. One can interpret this observation in the sense that defects in the primary metabolism, especially starch might be more severe for successful acclimation, highlighting the central role of starch within this process.

The deviations from the Col-0 sensitivity scores seem not to affect the already sensitive parts of the system around Frc and Glc, even though we see differences in the number of highly sensitive elements between different genotypes. This might be a safety feature of the compensation mechanism, to make sure that these sensitive vital parts are maintained against destabilization. Although, a substantial difference in subcellular reprogramming of sucrose cleavage by invertase between natural genotypes of *A. thaliana* has been demonstrated with model simulations (Weiszmann et al. 2018), this is not necessarily the case between mutant genotypes that are not specialized to grow in different habitats. The high control coefficients for Frc, Glc and neighboring reactions support the concept of a buffering effect of futile cycling against external perturbations (Atanasov et al. 2020; Brauner et al. 2015; Claeyssen et al. 2013; Geigenberger & Stitt 1991; Küstner et al. 2019; Nägele et al. 2010; Nägele et al. 2012), since it provides favorable conditions for a fine-tuned and flexible system. The fructokinase and glucokinase reactions also have a high sensitivity to changes in photosynthesis rates. Like this, the system can react fast and reliably to a sudden imbalance of sugars. The multiple feedback inhibitions provide a good backup, preventing overcompensation.

It is worth noticing that on day 0 (before acclimation), there is a significant quantitative difference between the deviations of the flavonol related and the starch related mutants for the rPGI and rSPS reactions. The *γ*_x_^nps^ values of the starch mutants are much closer to Col-0, while the absolute values of the flavonoid mutants are generally lower for these reactions. It might be the case that under normal conditions, these mutants naturally divert less carbon into the secondary pathways, since these are partially defective, so there is more carbon available that can potentially feed primary metabolism, rendering the initial steps less sensitive. This is in accordance with indications from the calibration, where the parameters k_ANT_ and k_r2_ have the lowest values for the two flavonol mutants.

Similar compensation programs, as indicated by the similar response patterns, for mutations in different metabolic branches hint to bi- or unidirectional signaling between these branches. Results from Kitashova et al. 2022 suggest a unidirectional signaling link between starch and flavonoid metabolism. These observations could possibly be connected.

A long-term objective for further investigation is the identification of potential common compensation mechanisms. It could be part of an already studied stress response. Additionally, the more downstream effects of the investigated mutations in combination with cold stress could be examined to see if and at which level the differences become more specific.

## Author Approval and Competing Interests

All authors have read and approved the manuscript.

The manuscript has not been accepted or published elsewhere.

There are no competing interests.

