## Supplemental Information for "Plant cold acclimation and its impact on sensitivity of carbohydrate metabolism"

### Supplementary Material

#### Section S1 – Metabolic Control Analysis

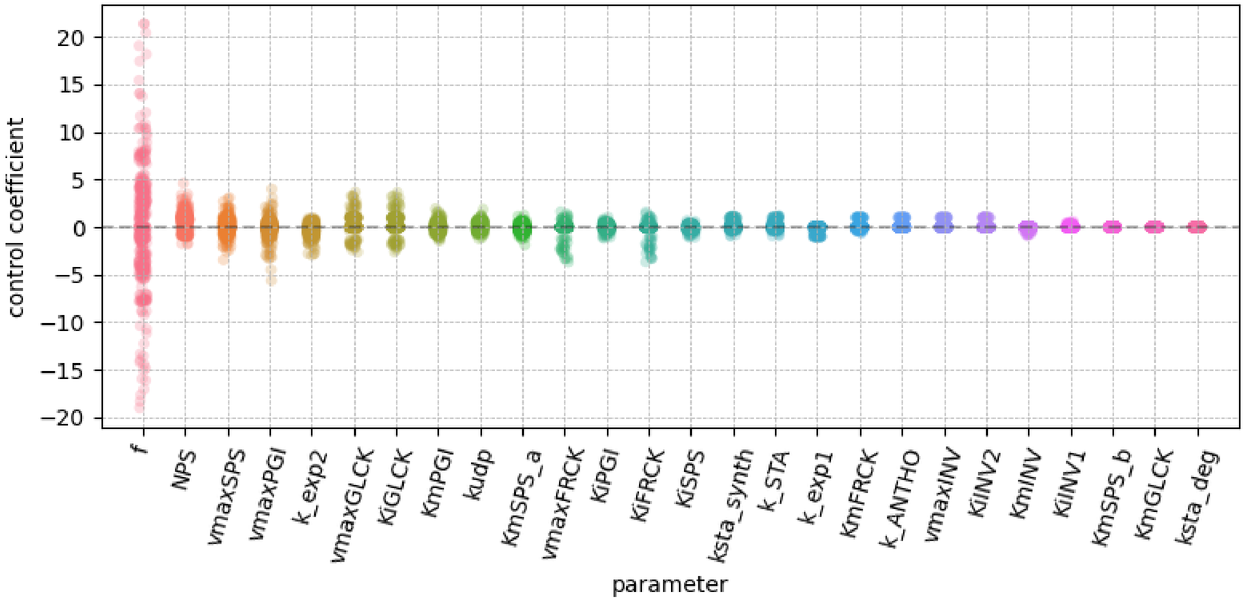


**Figure S1.1**: Control coefficient overview of all model parameters. Each dot represents the control coefficient of the respective parameter on either a flux or a concentration, for one genotype-day combination. This results in 5*5*15 = 375 control coefficients per parameter ( genotypes * days * (fluxes + components) ). The colors have no particular meaning and serve a better visual distinction between the columns.

For this analysis the control coefficients of various model parameters and fluxes were estimated and compared. These control coefficients quantify the relative change of a flux or concentration when another flux, concentration or a parameter is changed.

The equations read:

$$C_p^X = \frac{\mathrm{d}\ln X}{\mathrm{d}\ln p}$$

Where C is the control coefficient, X is the concentration or flux of interest and p is the parameter. Figure S1.1 clearly illustrates the strong influence of parameter *f* on the model components, compared to all other model parameters.

When looking at the control coefficients of *f* for each flux and concentration individually, one can notice a strong influence on Glc and Frc, which are both not directly connected to the *f*-dependent reactions. Also the influence on most other measures is relatively high compared to control coefficients of other parameters. It ranges to values of +/- 8 to 15 for at least one genotype-day combination, except for rNPS, which is completely independent.

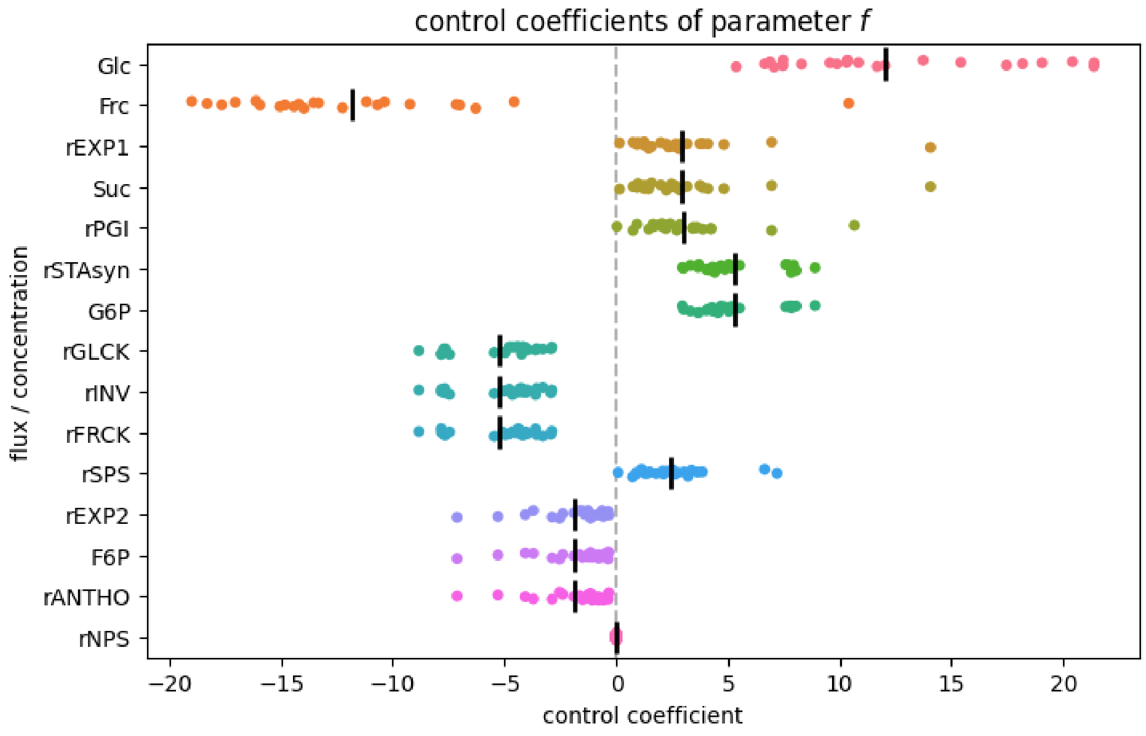


**Figure S1.2**: Control coefficients of parameter *f.* Each dot represents the control coefficient of *f* on the respective flux or concentration, for one genotype-day combination. This results in 5*5 = 25 control coefficients per measure ( genotypes * days ). The colors have no particular meaning and serve a better visual distinction between the columns.

### Section S2 – Principal component analysis

This section shows more results of the principal component analysis described in 3.3.2. Here, also the results of PC3 and PC4 are shown in Fig. S.2.1. One can notice that PC3 and PC4 each show a smaller spread along their axes, compared to PC1 and PC2. Also, when looking at the areas spanned by the respective values for each mutant or day, no particular pattern, like the clear separation of day 0 for PC1, can be observed.

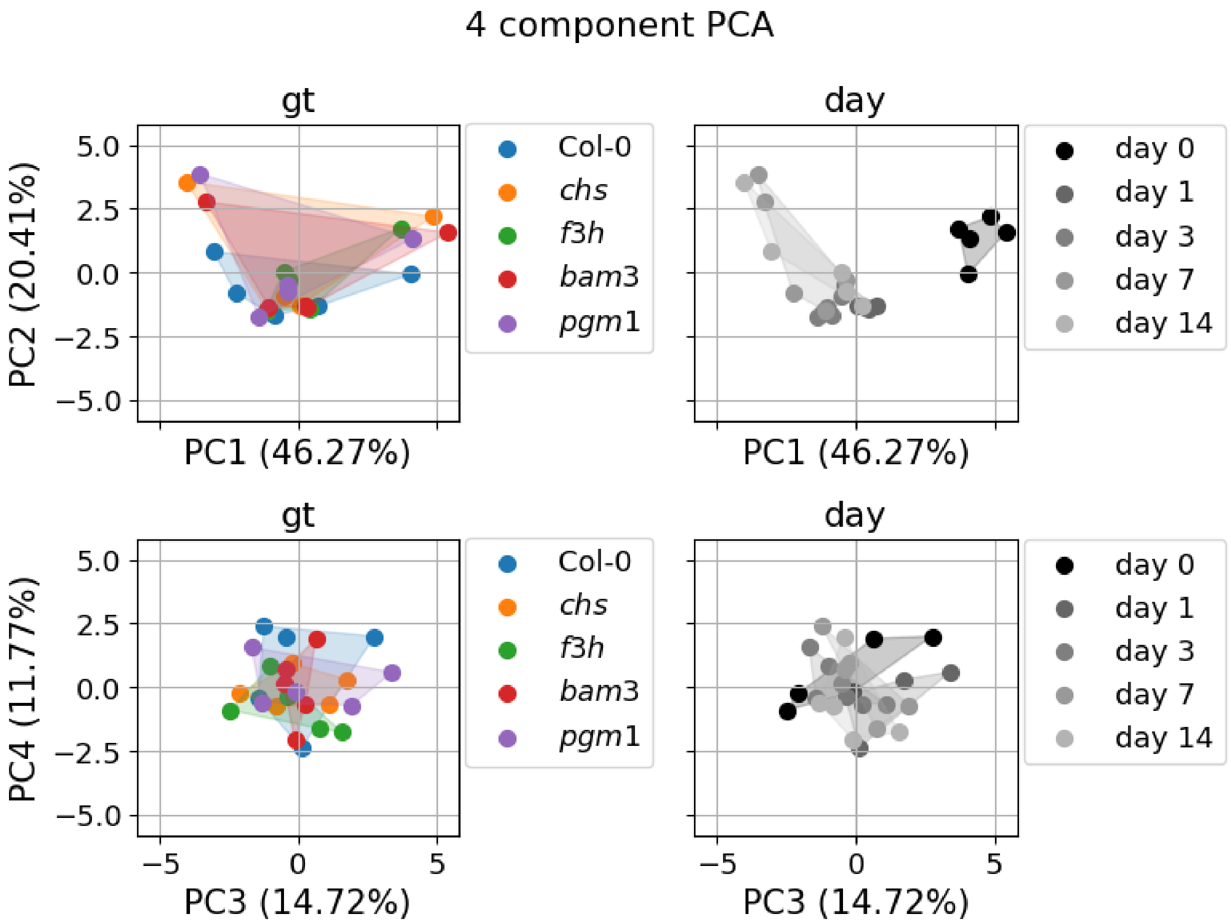


**Figure S2.1**: Four-component PCA. All panels show the results of a principle component analysis for variations in $\gamma_x^{nps}$. The upper two panels show PC1 against PC2, while the lower two panels show PC3 against PC4. The respective contribution of each component is given in % in the axis labels. In the left two panels, the data points of each mutant span an area and are color coded accordingly. The data points in the right two panels are the same, but span areas according to their days. Here, the corresponding grey scale saturation decreases for later days.
